## Supplementary Information for "A direct experimental test of Ohno’s hypothesis"

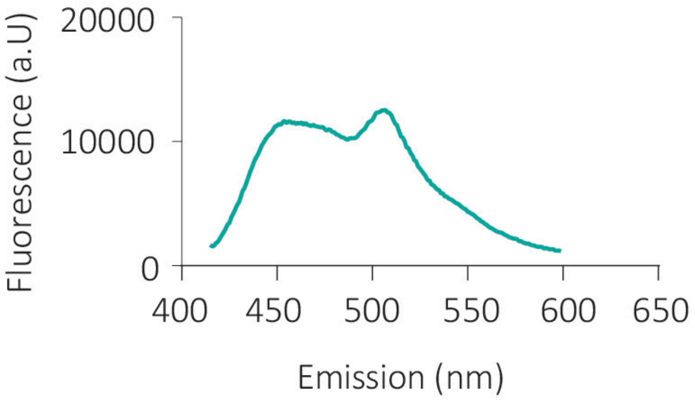


**Figure S1**. **Dual-color-emitting fluorescent protein coGFP**. Emission spectra of coGFP_S147G upon 388 nm excitation and pH 7. Two emission peaks at 456 nm (blue) and 507 nm (green) are detected.

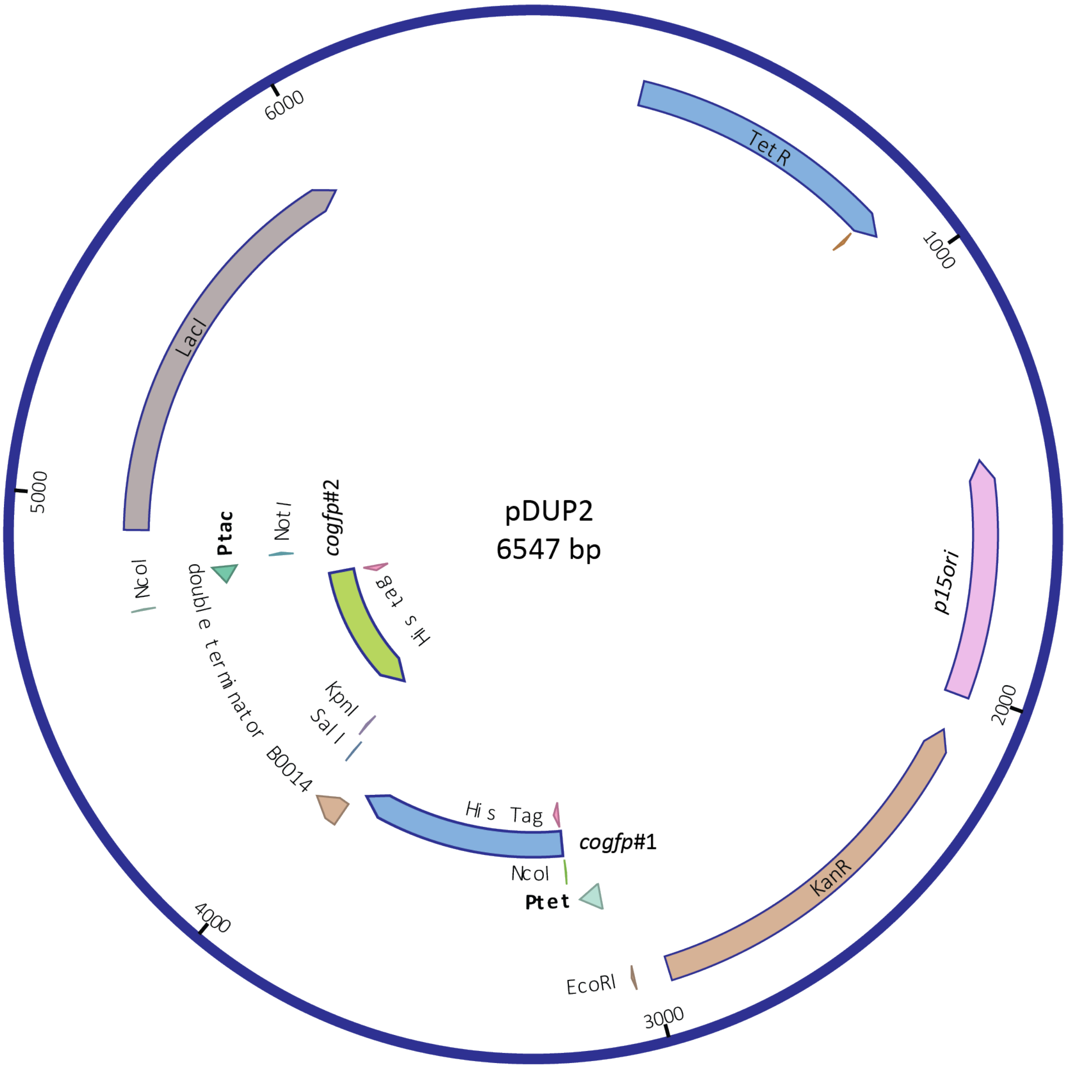


**Figure S2. Plasmid map of pDUP2 carrying the duplicated co*gfp* gene.** Double-copy of the *cogfp* gene, are facing each other, separated by a bidirectional terminator. The pDUP2 plasmid also includes the transcriptional repressors LacI and TetR and a kanamycin resistance gene. (#1: *cogfp* under the $P_{tet}$, #2: *cogfp* under the $P_{tac}$).

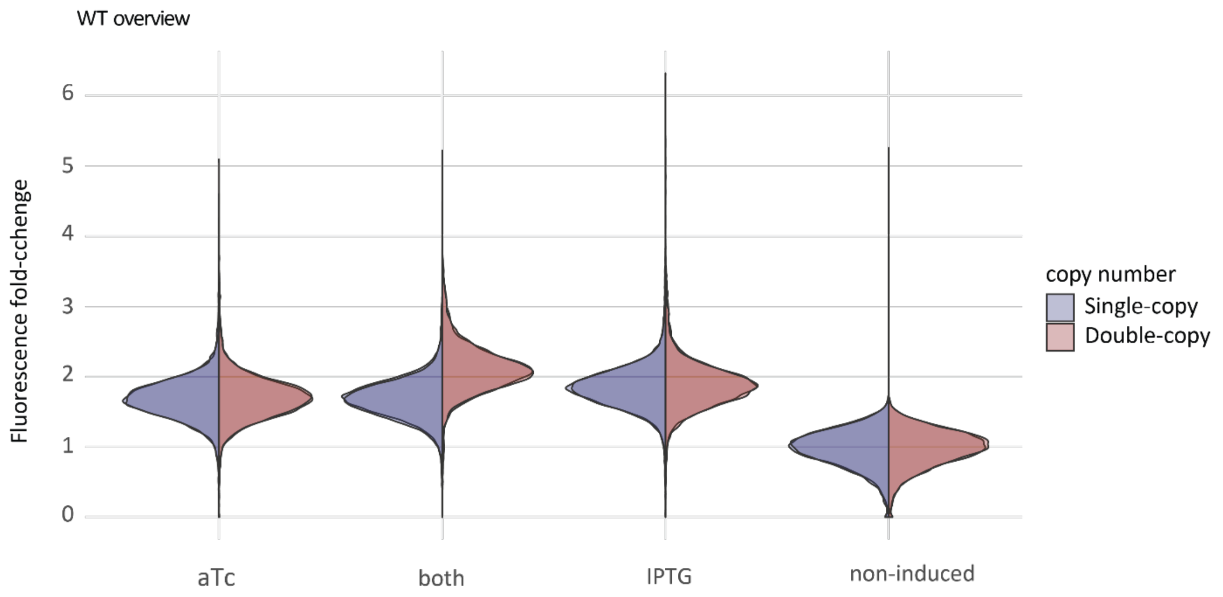


**Figure S3.** Expression levels of single-copy and double-copy constructs. Green fluorescence distribution of the ancestral single-copy (blue) and double-copy (red) populations measured by flow cytometry upon the induction with aTc, IPTG or both normalized to the non-induced controls.

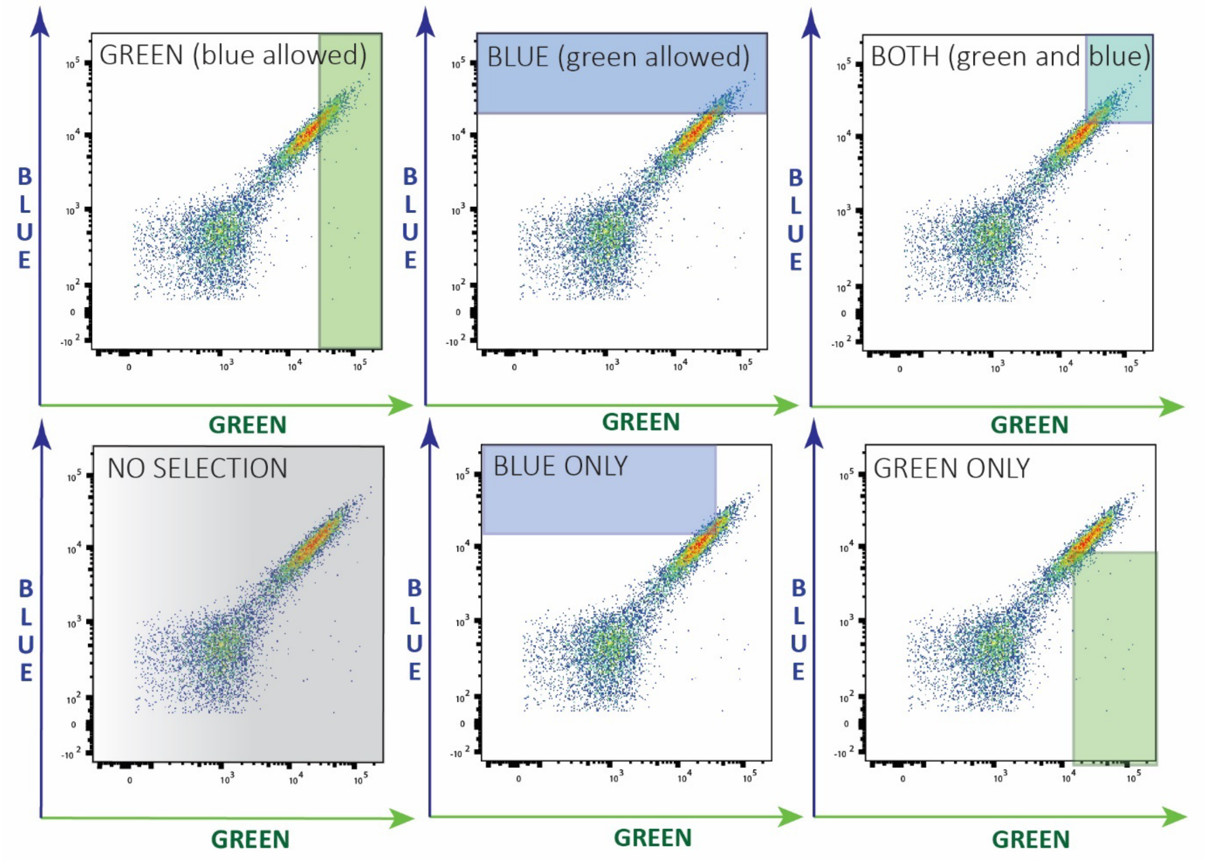


**Figure S4. Selection regimes based on cell's fluorescence phenotypes.** Flow cytometry plots of a first generation library after mutagenesis showing green (AmCyan) vs blue (DAPI) fluorescence. Highlighted regions indicate the gates used for the 6 different selection regimes: *green*: selection for green (no selection against blue fluorescence); *blue*: selection for blue (no selection against green fluorescence); *green and blue; green-only* (selection for green and against blue); *blue-only* (selection for blue and against green); and *no selection* for either fluorescence color.

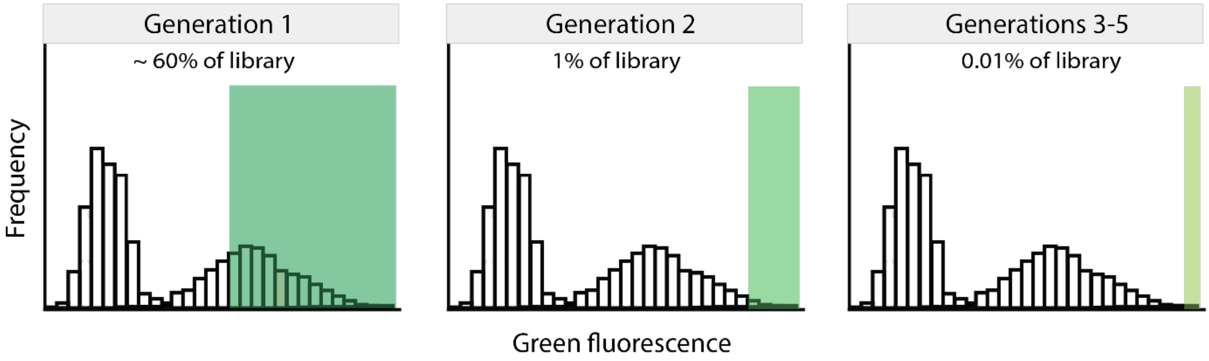


**Figure S5. Applied selection stringencies.** Histograms are schemes representing fluorescence distributions of the libraries. Single-copy wild type was used to set the selection threshold in the first generation of the evolution experiment (selected top 60%). In the upcoming generations, threshold is set on the mutant libraries (selected top 1% in the second and top 0.01% in the following rounds of the evolution experiment).

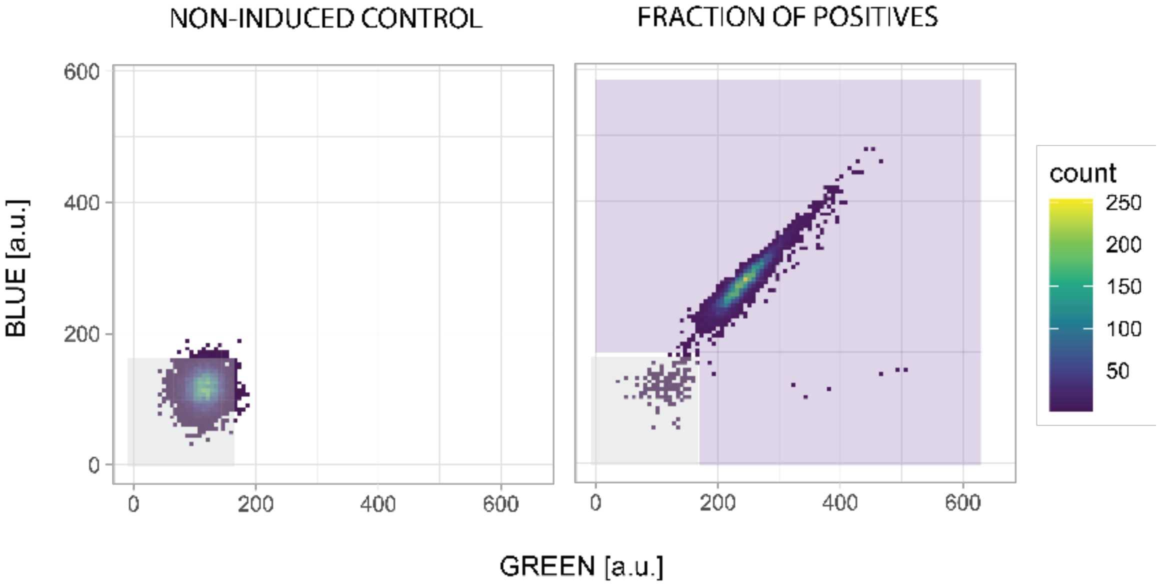


**Figure S6. Quantification of fluorescent cells.** Flow cytometry plots showing green (AmCyan) vs blue (DAPI) fluorescence. A non-induced control was used to set the gate (grey). Cells outside this gate (violet area) are considered fluorescent.

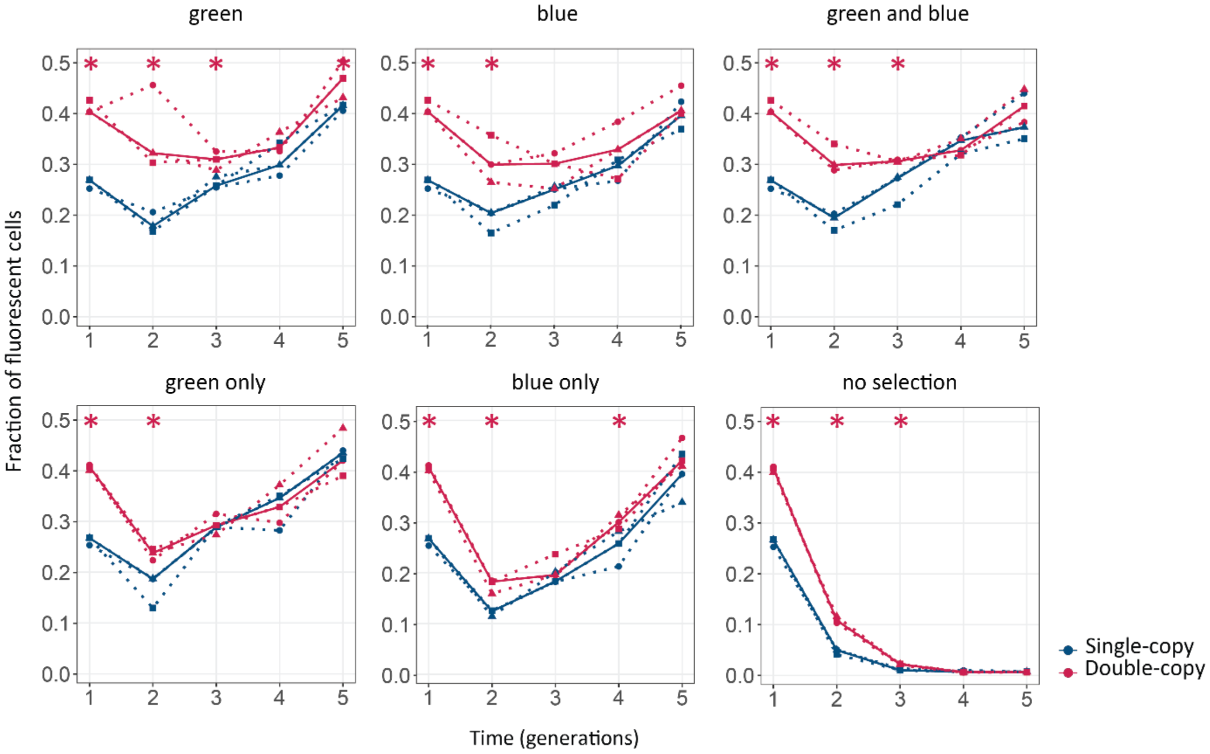


**Figure S7. Gene duplication increases mutational robustness.** The vertical axis shows mutational robustness, measured as the percentage of cells that maintain their fluorescence after mutagenesis, as a function of time (in generations of directed evolution) on the horizontal axis. Thick blue and red lines stand for the median fraction of fluorescent cells for single-copy and double-copy mutant libraries respectively, while dotted lines and circles indicate data from the three biological replicates. The corresponding selection regime is indicated at the top. One tailed Mann-Whitney tests, * p≤0.05, n=3.

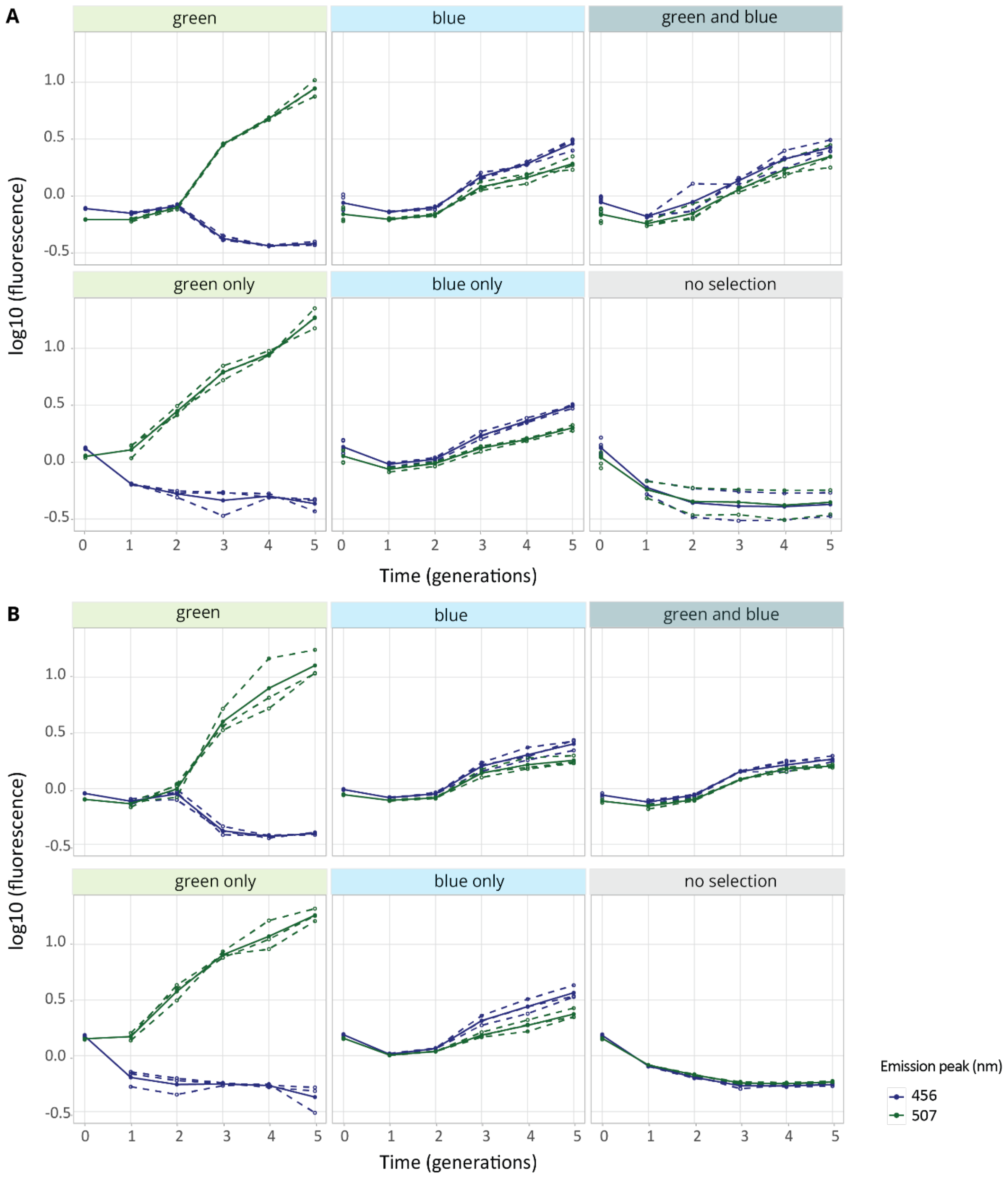


**Figure S8. Fluorescence levels during evolution experiment.** Fluorescence levels of the (**A**) single- and (**B**) double-copy populations evolved under the indicated selection regimes throughout five generations of the evolution (1-5). Shown is the fluorescence (log10) at blue (456 nm) and green (507 nm) emission peaks upon excitation at 388 nm normalized to the ancestral population. Thick blue and green lines show the mean fluorescence of the populations, while dotted lines indicate data from the three biological replicates.

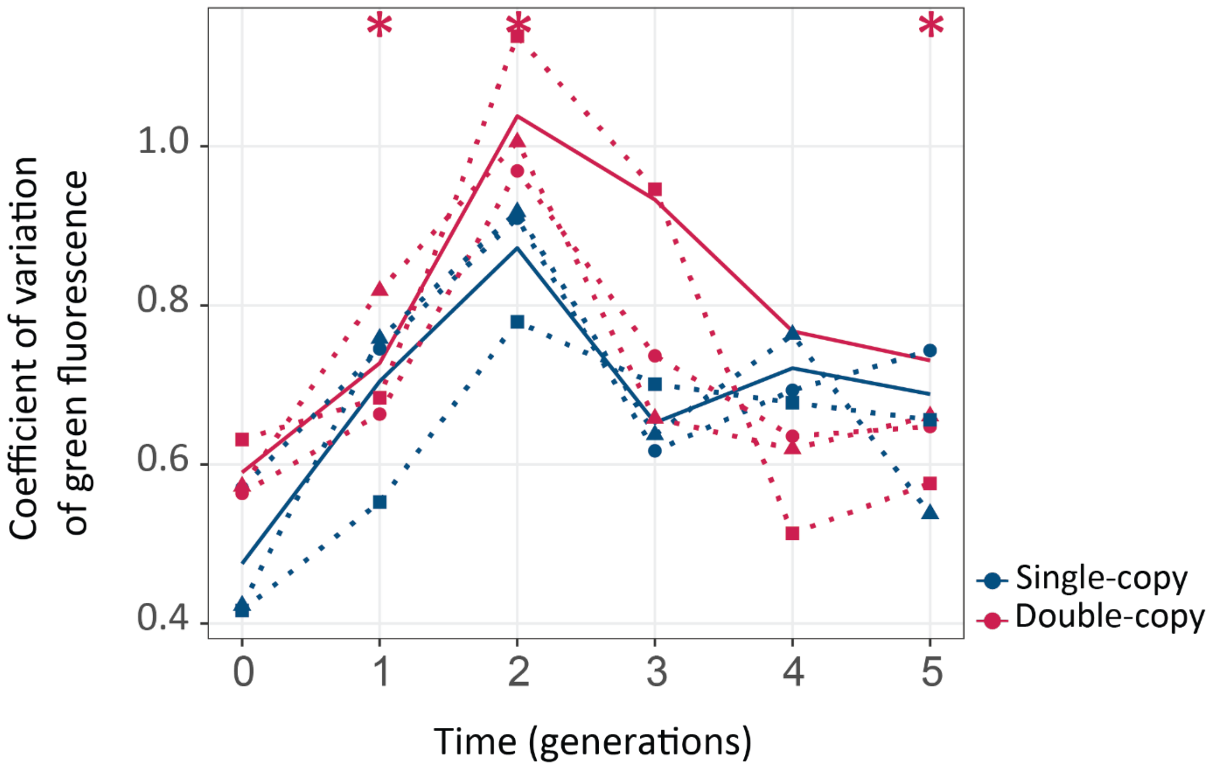


**Figure S9**. **Coefficient of variation of fluorescence**. Populations evolved under selection for green. The vertical axis shows the coefficient of variation (CV) of fluorescence (standard deviation/mean) as a function of time (generations) on the horizontal axis. Thick blue and red lines stand for median variance for single or double-copy populations, respectively, while dotted lines indicate data from the three biological replicates. One tailed Mann-Whitney tests, * $p<0.05$, $n=3$.

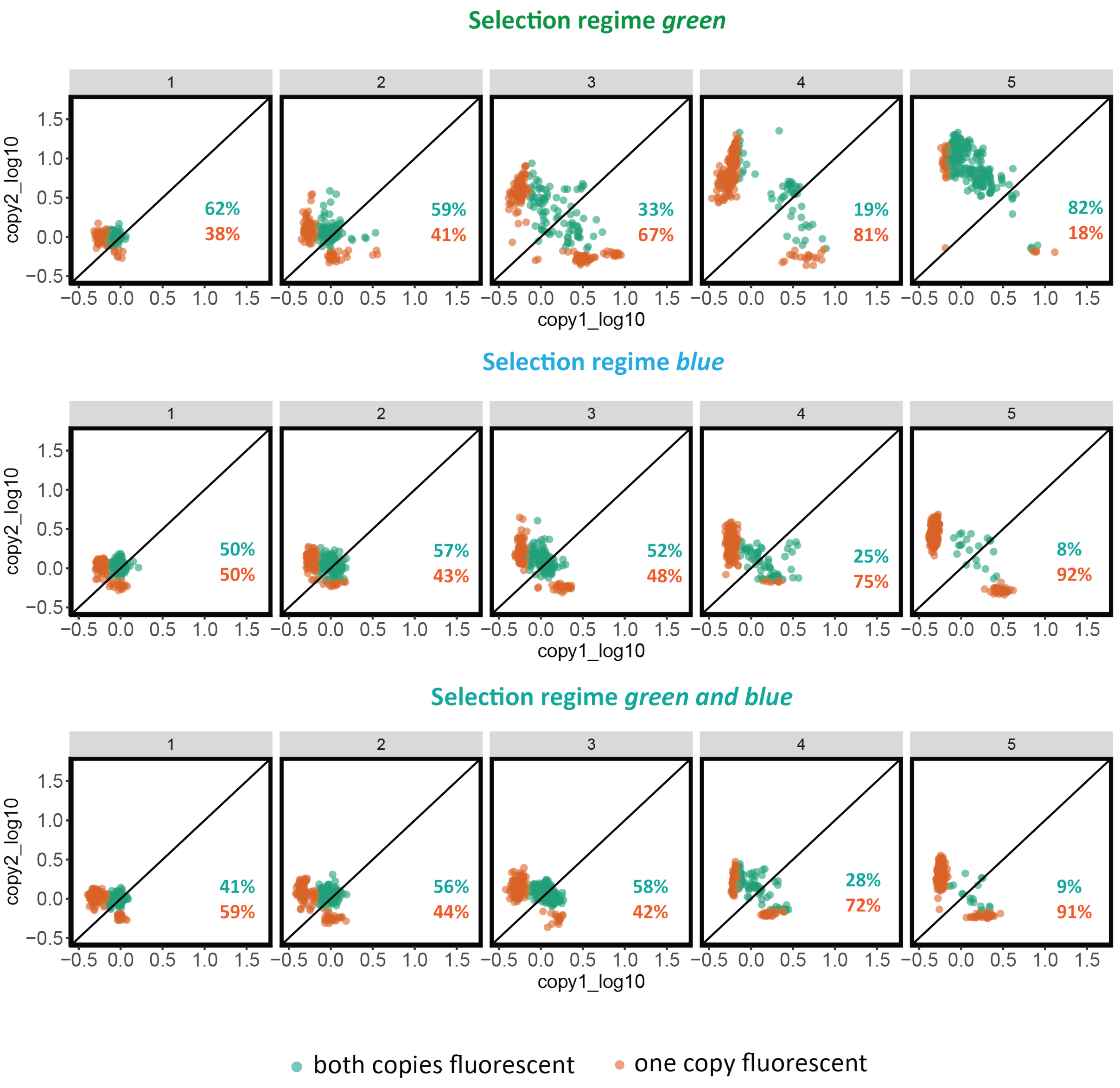


**Figure S10. Copy maintenance vs copy loss.** Green fluorescence (for the regimes "green" and "green and blue") and blue fluorescence (for the regime "blue"), from copy 1 (x-axes) and copy 2 (y-axes) of the double-copy populations, normalized to the fluorescence of the ancestral construct (0/0), displayed on log axes. Color code indicates if one (green) or two (orange) copies are still displaying fluorescence. Generations are indicated at the top. Data points represent mean from two technical replicates.

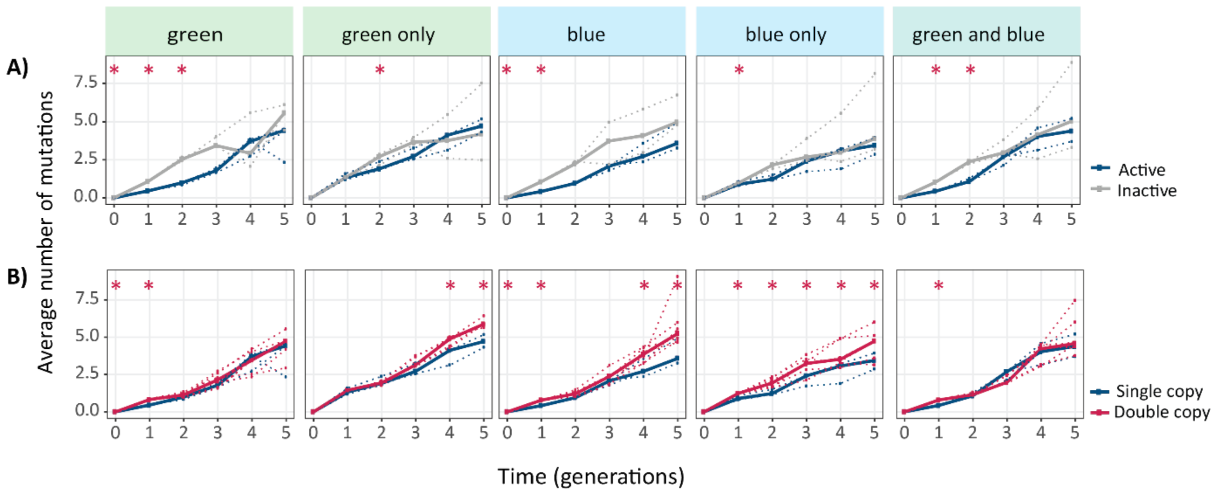


**Figure S11**. **Average number of mutations** for single- and double-copy populations, evolved under the indicated selection regimes. The vertical axes show the average number of non-synonymous mutations per *cogfp* gene, as a function of time (in generations of directed evolution) on the horizontal axes. A) Average number of mutations per coGFP in single-copy populations (blue: active copy, grey: inactive copy), B) average number of mutations per coGFP of the active copy in single-copy populations (blue) vs both copies in double-copy populations (pink). Thick lines stand for the median of populations, while dotted lines and circles indicate data from the three biological replicates., Mann-Whitney tests, * $p<0.05$, $n=3$.

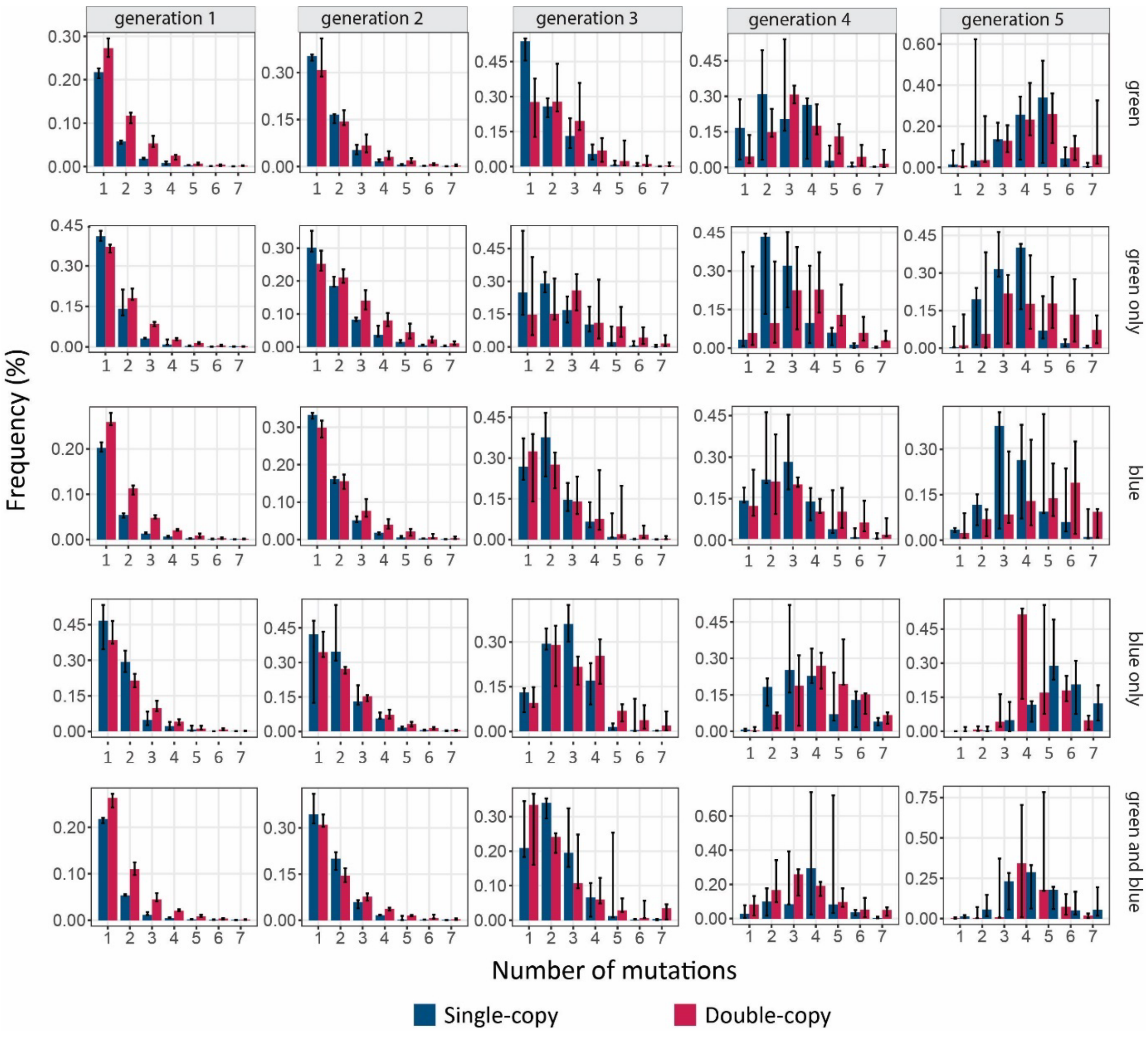


**Figure S12. Frequency distribution of the number of mutations.** The vertical axes show the frequency of mutations, as a function of the number of mutations per *cogfp* gene on the horizontal axes. Blue and red bars stand for single-copy and double-copy populations, respectively. Generations of directed evolution are indicated at the top and selection regimes are indicated on the right. Average and standard deviation of three biological replicates.

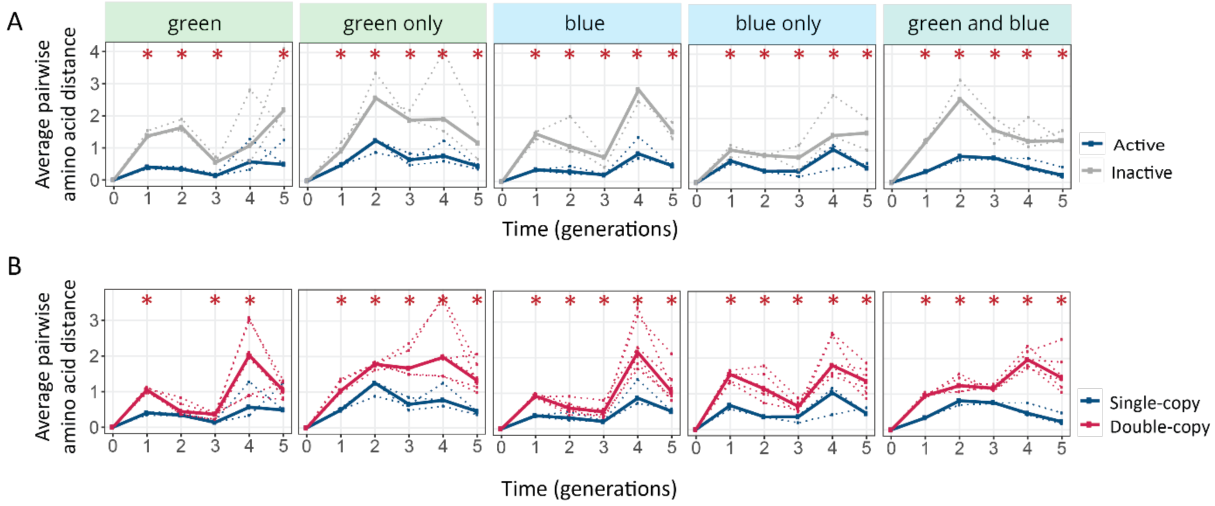


**Figure S13. Populations with two gene copies are showing increased genetic diversity.** The horizontal axes of all panels show time in generations for the indicated selection regimes. **A)** Average pairwise amino acid distance for coGFP molecules in single-copy populations (blue: active copy, grey: inactive copy), **B)** Average pairwise amino acid distance for coGFP molecules of the active copy in single-copy populations (blue) vs both copies in double-copy populations (pink). One tailed Mann-Whitney tests, * $p<0.05$, $n=3$.

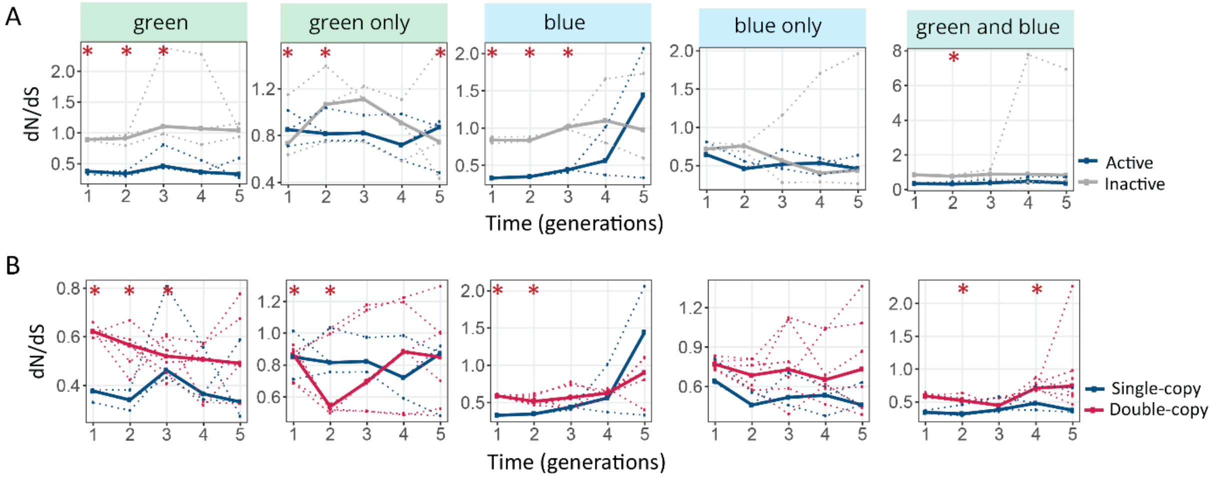


**Figure S14. Populations with two gene copies are showing higher dN/dS ratios. A)** dN/dS ratio in single-copy populations (blue: active copy, grey: inactive copy), **B)** dN/dS ratio of the active copy in single-copy populations (blue) vs both copies in double-copy populations (pink). Thick lines represent the median over three replicate populations, while dotted lines indicate data from the individual biological replicates. One tailed Mann-Whitney tests, * $p \leq 0.05$, $n=3$.

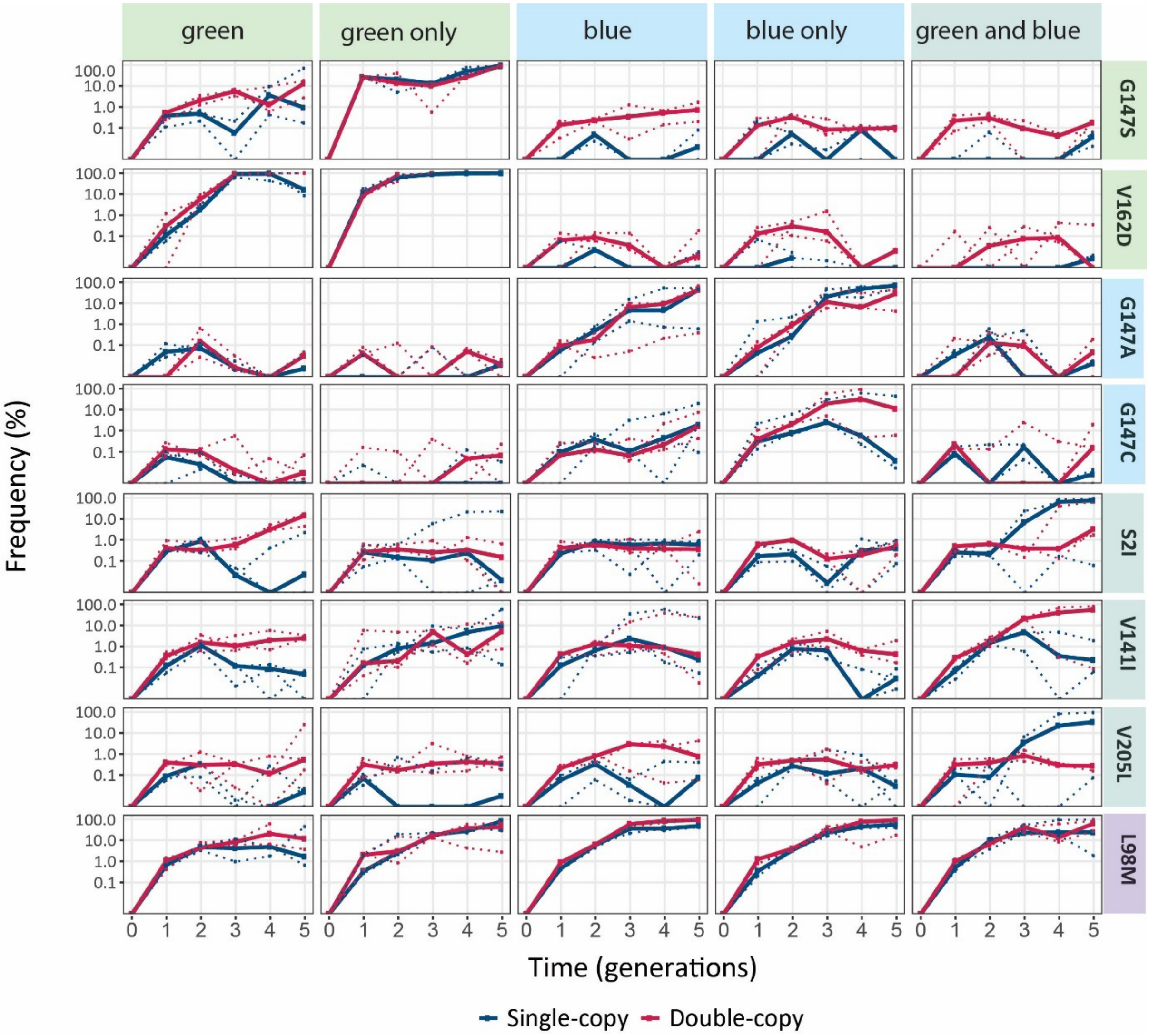


**Figure S15. Enriched mutations.** The vertical axes show the frequency of indicated mutations in the populations under indicated selection regimes, as a function of time (in generations of directed evolution) on the horizontal axis. Thick blue and red lines stand for the median frequencies for single-copy and double-copy populations, respectively, while dotted lines and circles indicate data from the three biological replicates.

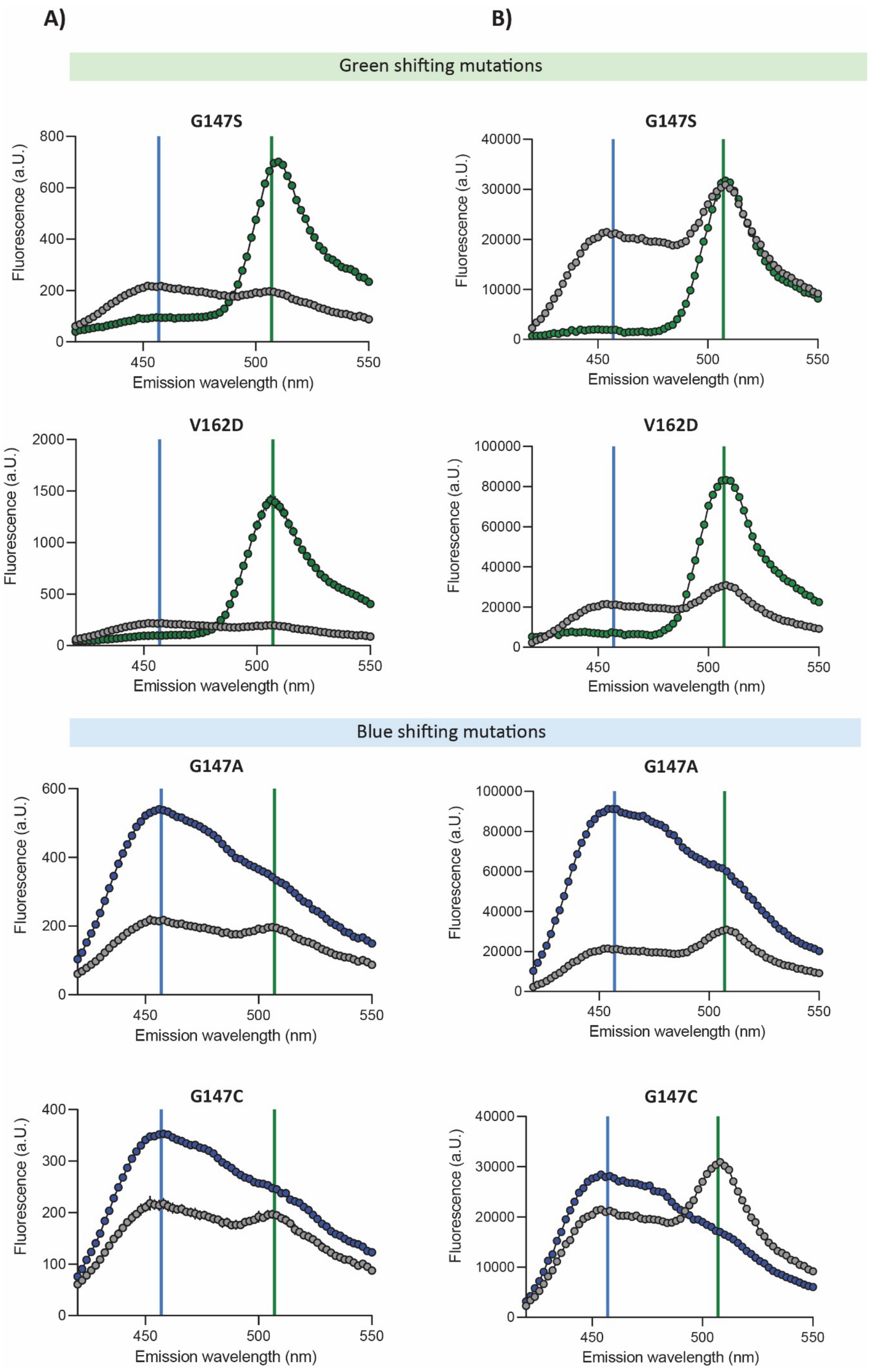

A)
B)
Green shifting mutations
G147S
Fluorescence (a.U.)
Emission wavelength (nm)
V162D
Blue shifting mutations
G147A
G147C

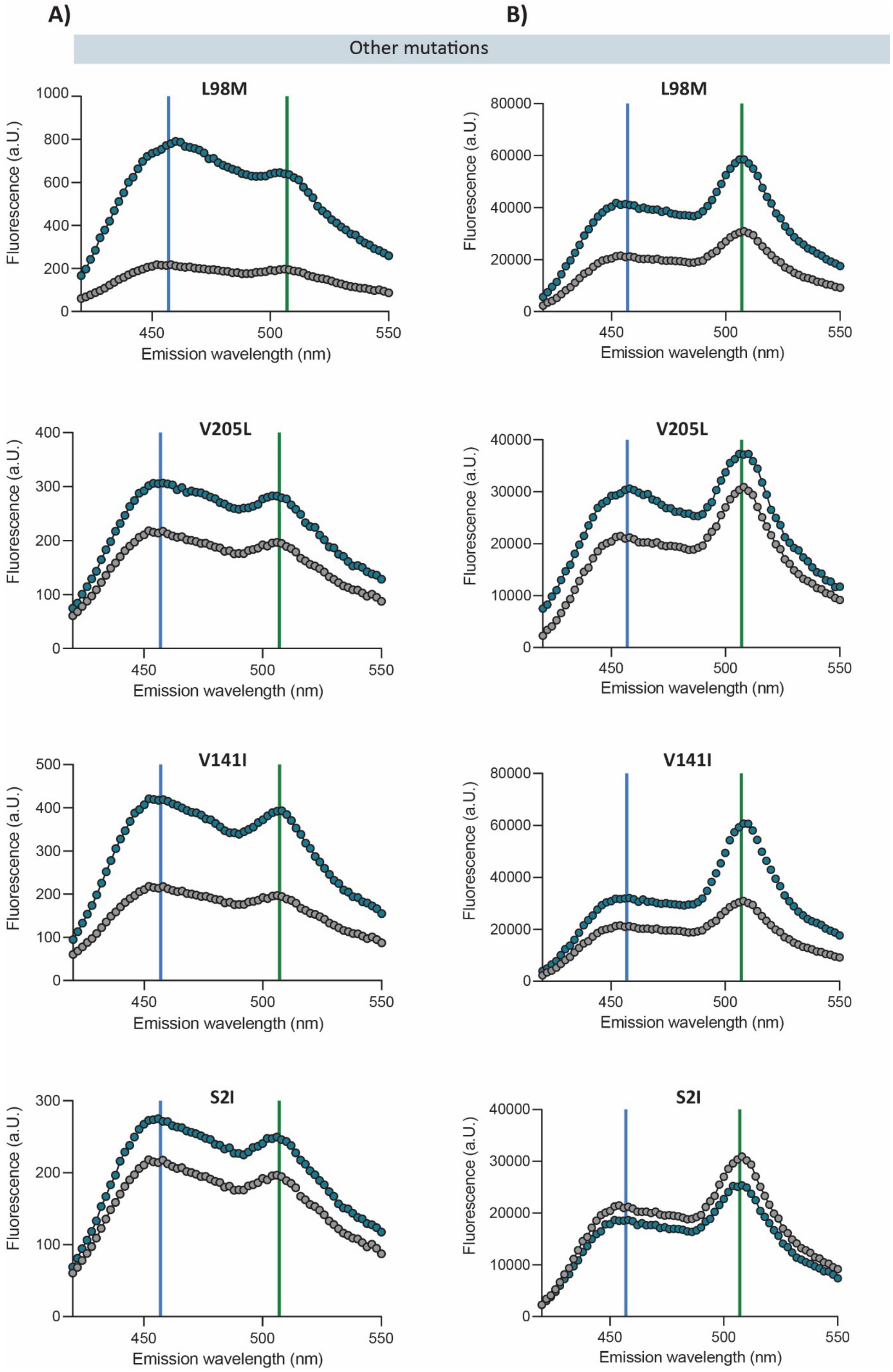


**Figure S16. Emission spectra of engineered coGFP variants.** The vertical axes show the fluorescence intensities as a function of the emission wavelengths upon 388 nm excitation of the indicated coGFP variants. Vertical lines indicate blue (456 nm) and green (507 nm) emission peaks. In color, the engineered variants compared to the ancestral protein in grey. **A)** Fluorescence of bacterial cells expressing a single copy of the indicated coGFP variant. **B)** Fluorescence of purified protein of the indicated coGFP variant (0.05 mg/ml).

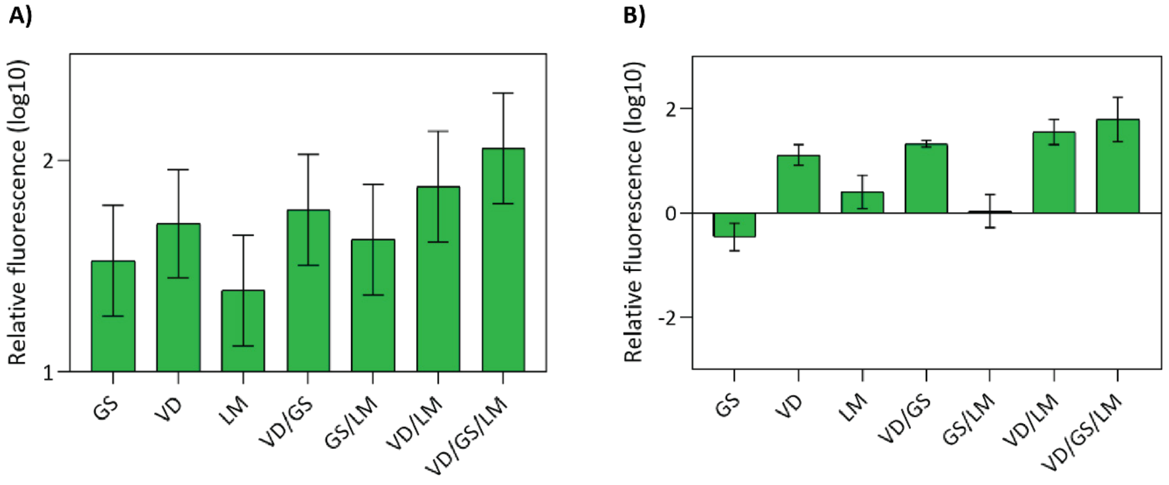


**Figure S17. Relative green fluorescence of the engineered variants compared to the ancestral variant.** The vertical axes show the green (507 nm) fluorescence intensities (log10) of the indicated variant relative to the that of the ancestral variant upon excitation at 388 nm. **A)** Fluorescence of bacterial cells expressing a single copy of the indicated coGFP variant. **B)** Fluorescence of purified coGFP protein after size-exclusion chromatography to remove unfolded protein. Tested variants: G147S (GS), V162D (VD), L98M (LM), G147S/V162D (GS/VD), G147S/L98M (GS/LM), V162D/L98M (VD/LM), G147S/V162D/L98M (GS/VD/LM)

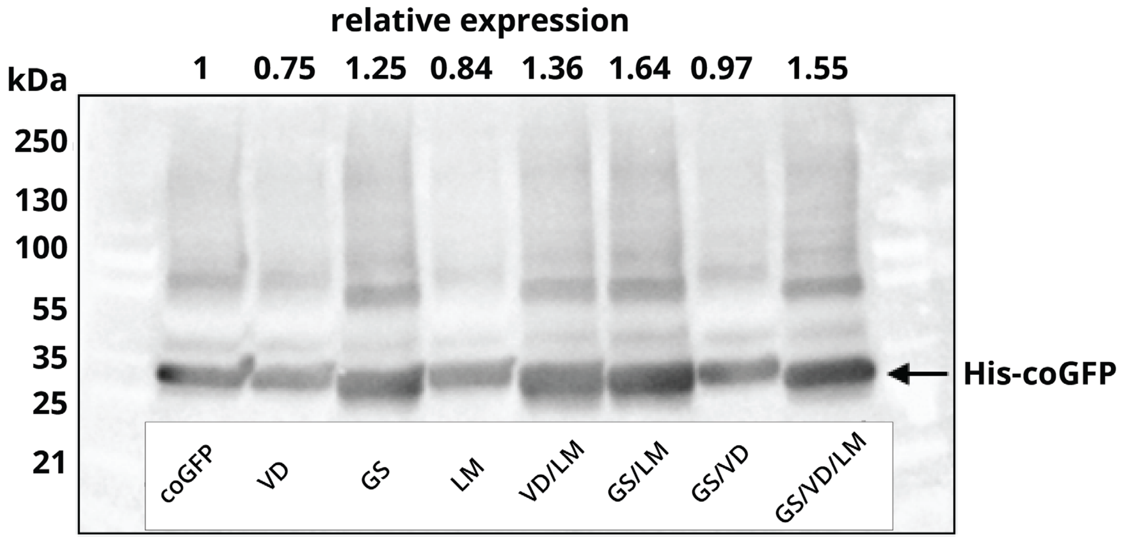


**Figure S18. Expression levels of the engineered coGFP variants.** Western blot of the soluble cell lysate fraction using a primary mouse antibody against the His-tag on coGFP and a secondary goat anti-mouse antibody conjugated to horseradish peroxidase (HRP) for chemiluminescent detection. Relative expression levels compared to the ancestral variant coGFP 147G (wt) are indicated at the top. Tested variants: coGFP (wt), G147S (GS), V162D (VD), L98M (LM), G147S/V162D (GS/VD), G147S/L98M (GS/LM), V162D/L98M (VD/LM), G147S/V162D/L98M (GS/VD/LM).

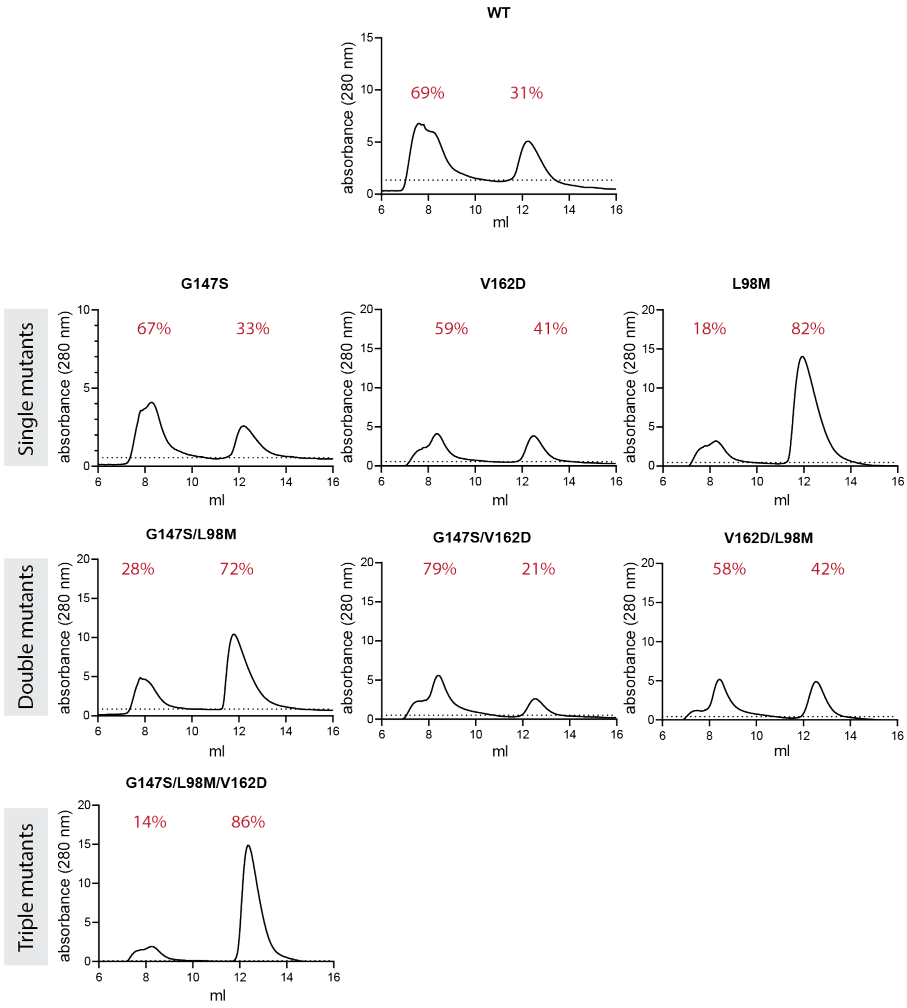


**Figure S19. Analysis of fractions of folded protein.** Purified coGFP proteins were run on a size-exclusion column. The vertical axes show the absorbance (280 nm) - a proxy for protein concentration as a function of the elution volume (in ml). Two main peaks are detected: The first one (at ~8ml) corresponds to aggregated unfolded proteins. The second one (at ~13ml) corresponds to folded protein. Fractions of unfolded and folded protein are indicated in red numbers above the peaks.

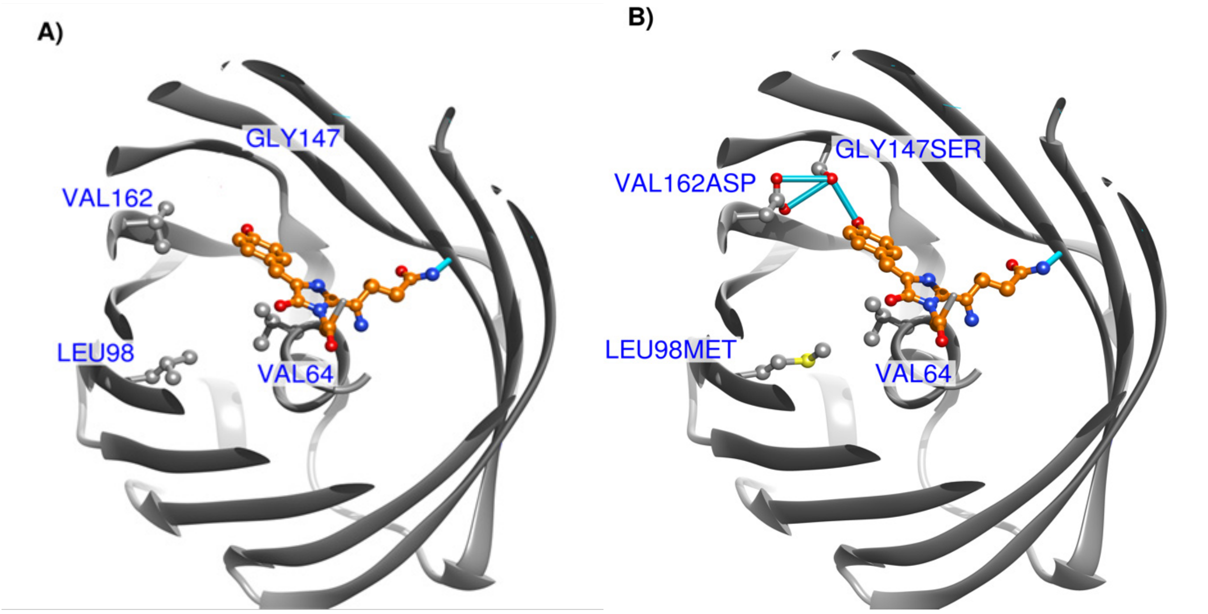


**Figure S20. Structure of coGFP. A)** Homology model of the ancestral coGFP with positions affected by key mutations presented in panel B), top view. **B)** Homology model of the coGFP triple mutant L98M, G147S, V162D chromophore region, top view. Top part of the protein is made invisible for clarity. Grey: coGFP in ribbon representation and carbon atoms, orange: mature chromophore; red: oxygen atoms; blue: nitrogen atoms; yellow: sulfur atoms; bright blue: hydrogen bonds.

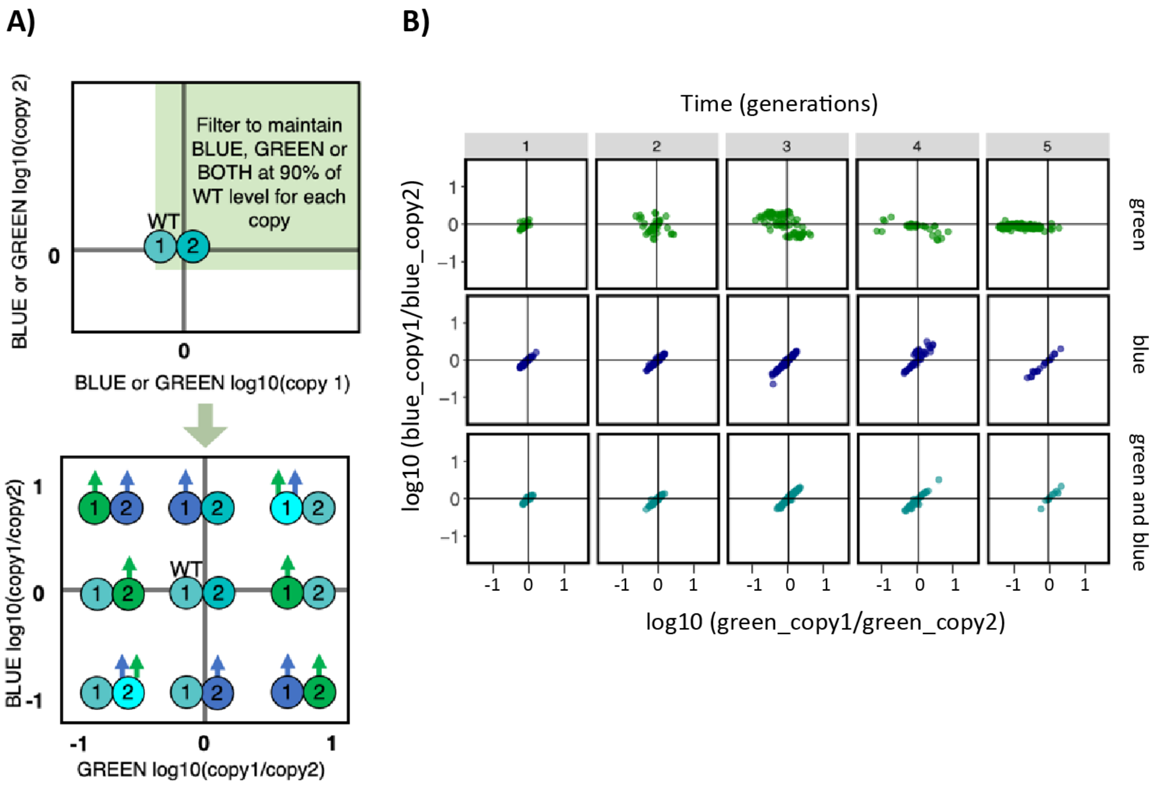


**Figure S21. Analysis of variants with two active gene copies. A)** Scheme to explain the data analysis. Both copies were individually induced, and blue and green fluorescence were measured. This analysis only looks at variants where both copies have green or blue fluorescence of at least 90% of the ancestral variant. For those we plotted the ratio of copy1/copy2 for blue (x-axis) and green (y-axis) fluorescence (log10 scale). The bottom scheme explains where the different scenarios will lie. For example, data points in the upper left corner have copy 1 improved in green and copy 2 improved in blue compared to the ancestral (WT) protein. Data points in the upper right corner have copy 1 improved in blue and green fluorescence, while copy 2 did not change much. **B)** Actual data as explained in A). We did not find cases where one copy improves in green and the other copy improves in blue.

**Table S1. Summary of SMRT sequencing results**. Number of reads sequenced by SMRT sequencing and mean number of amino-acid changes per *cogfp* gene. X, Y, Z: replicate populations. 1-5: generations of evolution.

**Single-copy populations**

| | mean number of amino acid changes | | number of reads | mean number of amino acid changes | | number of reads | mean number of amino acid changes | | number of reads |
|---|---|---|---|---|---|---|---|---|---|
| Starting Libraries | X | 1.989 | 3379 | Y | 1.900 | 707 | Z | 1.869 | 4610 |
| GREEN | X active copy | X inactive copy | | Y active copy | Y inactive copy | | Z active copy | Z inactive copy | |
| 1 | 0.434 | 1,080 | 3378 | 0.473 | 1,052 | 3585 | 0.458 | 1,102 | 2058 |
| 2 | 0.838 | 2,423 | 13241 | 0.994 | 2,540 | 25021 | 1,037 | 2,675 | 15072 |
| 3 | 1,649 | 4,006 | 23561 | 1,944 | 3,422 | 23204 | 1,789 | 3,386 | 12514 |
| 4 | 2,738 | 5,576 | 1445 | 3,853 | 2,926 | 2205 | 3,716 | 2,062 | 279 |
| 5 | 4,428 | 6,109 | 6316 | 4,503 | 4,451 | 1090 | 2,343 | 5,577 | 2420 |
| GREEN ONLY | | | | | | | | | |
| 1 | 1,189 | 1,312 | 1869 | 1,592 | 1,367 | 4324 | 1,325 | 1,316 | 3041 |
| 2 | 1,806 | 2,861 | 2028 | 2,381 | 2,150 | 955 | 1,911 | 2,715 | 1607 |
| 3 | 2,720 | 3,957 | 940 | 3,257 | 3,509 | 805 | 2,550 | 3,659 | 1262 |
| 4 | 4,139 | 5,452 | 820 | 4,122 | 3,764 | 1606 | 3,138 | 2,593 | 904 |
| 5 | 5,176 | 7,513 | 5986 | 4,715 | 4,201 | 9848 | 4,318 | 2,480 | 8358 |
| BLUE | | | | | | | | | |
| 1 | 0.428 | 1,053 | 3183 | 0.383 | 1,021 | 994 | 0.417 | 1,067 | 1730 |
| 2 | 0.954 | 2,234 | 25768 | 0.926 | 2,164 | 8096 | 0.973 | 2,375 | 21774 |
| 3 | 1,787 | 4,960 | 21638 | 2,080 | 3,731 | 26241 | 2,161 | 2,204 | 29250 |
| 4 | 2,720 | 5,815 | 1100 | 3,573 | 4,084 | 1122 | 2,362 | 2,979 | 1152 |
| 5 | 3,584 | 6,742 | 10493 | 4,919 | 4,973 | 16071 | 3,267 | 4,794 | 5743 |
| BLUE ONLY | | | | | | | | | |
| 1 | 1,077 | 0.941 | 2380 | 0.893 | 1,000 | 2287 | 0.807 | 0.987 | 2943 |
| 2 | 1,504 | 2,249 | 32035 | 1,231 | 2,167 | 33968 | 1,160 | 1,910 | 35922 |
| 3 | 2,456 | 2,680 | 14421 | 2,419 | 3,890 | 9838 | 1,728 | 2,451 | 24708 |
| 4 | 3,068 | 2,391 | 948 | 3,225 | 5,563 | 1005 | 1,896 | 2,996 | 2186 |
| 5 | 3,447 | 3,193 | 10725 | 3,935 | 8,152 | 10294 | 2,864 | 3,899 | 5941 |
| GREEN AND BLUE | | | | | | | | | |
| 1 | 0.431 | 1,032 | 2827 | 0.435 | 1,009 | 3118 | 0.41 | 1,025 | 3672 |
| 2 | 1,078 | 2,396 | 1290 | 1,293 | 2,236 | 1168 | 1,005 | 2,386 | 1543 |
| 3 | 2,124 | 2,960 | 1656 | 2,695 | 2,830 | 2049 | 2,714 | 3,818 | 1195 |
| 4 | 4,029 | 2,571 | 1129 | 3,139 | 4,138 | 2132 | 4,575 | 5,848 | 1243 |
| 5 | 4,389 | 3,307 | 7124 | 3,706 | 5,033 | 11701 | 5,209 | 8,884 | 8003 |

### Double-copy populations

| | mean number of amino acid changes | | number of reads | mean number of amino acid changes | | number of reads | mean number of amino acid changes | | number of reads |
|---|---|---|---|---|---|---|---|---|---|
| Starting Libraries | X | 2.009 | 1494 | Y | 1.997 | 4061 | Z | 2.058 | 1083 |
| GREEN | X copy 1 | X copy 2 | | Y copy 1 | Y copy 2 | | Z copy 1 | Z copy 2 | |
| 1 | 0.83 | 0.778 | 765 | 0.808 | 0.826 | 1685 | 0.872 | 0.848 | 749 |
| 2 | 0.978 | 1,062 | 11012 | 1,270 | 1,127 | 18784 | 1,344 | 1,151 | 11933 |
| 3 | 1,582 | 1,646 | 22785 | 2,741 | 2,602 | 15681 | 2,296 | 2,001 | 11697 |
| 4 | 3,456 | 3,481 | 2016 | 4,034 | 4,208 | 870 | 2,526 | 2,349 | 2166 |
| 5 | 4,708 | 4,755 | 19201 | 4,793 | 5,542 | 8542 | 2,934 | 4,193 | 10122 |
| GREEN ONLY | | | | | | | | | |
| 1 | 1,488 | 1,456 | 2754 | 1,440 | 1,460 | 2553 | 1,421 | 1,412 | 1278 |
| 2 | 1,770 | 1,879 | 861 | 1,915 | 1,978 | 2442 | 2,076 | 2,057 | 3016 |
| 3 | 3,185 | 3,112 | 720 | 3,121 | 3,043 | 769 | 3,687 | 3,766 | 1172 |
| 4 | 4,987 | 5,024 | 2860 | 4,418 | 4,295 | 1937 | 4,956 | 4,819 | 2130 |
| 5 | 5,876 | 5,649 | 9495 | 5,888 | 5,897 | 4528 | 6,424 | 5,762 | 16576 |
| BLUE | | | | | | | | | |
| 1 | 0.791 | 0.77 | 3143 | 0.768 | 0.781 | 1451 | 0.825 | 0.826 | 1905 |
| 2 | 1,359 | 1,052 | 27650 | 1,485 | 1,103 | 10375 | 1,329 | 1,011 | 13705 |
| 3 | 2,421 | 1,810 | 13949 | 3,113 | 2,341 | 16774 | 2,447 | 1,873 | 16611 |
| 4 | 3,673 | 4,053 | 1381 | 4,360 | 4,286 | 2428 | 3,229 | 3,304 | 3389 |
| 5 | 9,067 | 4,654 | 7046 | 5,999 | 4,949 | 12312 | 5,550 | 4,780 | 16739 |
| BLUE ONLY | | | | | | | | | |
| 1 | 1,218 | 1,185 | 2771 | 1,261 | 1,211 | 1263 | 1,277 | 1,297 | 1536 |
| 2 | 2,146 | 1,644 | 27424 | 2,346 | 1,728 | 14607 | 2,285 | 1,666 | 14189 |
| 3 | 3,149 | 2,124 | 11041 | 3,529 | 2,822 | 11402 | 3,837 | 3,339 | 10382 |
| 4 | 3,637 | 3,280 | 3063 | 3,408 | 2,969 | 1577 | 4,907 | 4,936 | 2177 |
| 5 | 4,765 | 3,136 | 10307 | 4,712 | 3,615 | 4924 | 6,016 | 5,109 | 8487 |
| GREEN AND BLUE | | | | | | | | | |
| 1 | 0.792 | 0.828 | 1819 | 0.801 | 0.818 | 1283 | 0.755 | 0.78 | 1430 |
| 2 | 1,309 | 1,286 | 1004 | 1,108 | 1,141 | 2907 | 1,113 | 1,156 | 2318 |
| 3 | 1,914 | 1,986 | 1347 | 1,979 | 1,983 | 1071 | 2,433 | 2,462 | 790 |
| 4 | 3,080 | 3,174 | 1925 | 4,524 | 4,449 | 2395 | 4,199 | 4,259 | 2600 |
| 5 | 4,495 | 3,773 | 8131 | 7,478 | 4,660 | 9477 | 6,021 | 4,349 | 6450 |

**Table S2.** Values of structural stability difference DDG between homology models of coGFP structures and the mutated structures, calculated with Buildmodel FoldX function. The results pointing to destabilizing effects are colored orange and stabilizing or synergistic are colored green.

| mutation | ΔΔG by FoldX (kcal/mol) | ∑ of individual single mutant ΔΔG (kcal/mol) | difference |
|---|---|---|---|
| **L98M** | -0.25 | -0.25 | 0 |
| **G147S** | -0.57 | -0.57 | 0 |
| **V162D** | +3.46 | +3.46 | 0 |
| **G147S_V162D** | +3.33 | +2.86 | + 0.47 |
| **L98M_G147S_V162D** | +1.95 | +2.64 | -0.69 |
| **L98M_V162D** | +3.28 | +3.21 | 0.08 |
| **L98M_G147S** | -1.05 | -0.82 | -0.23 |

**Supplementary notes to Table S2.** The G147S mutation alone is stabilizing the structure because of the hydrogen bond that it is forming with the chromophore Y75 residue (**Table S2**). The mutation of L98M is also changing favorably the structural stability due to longer Met side chain, that is able to form stabilizing hydrophobic interactions with V64 residue of the central helix of coGFP (**Table S2, Figure S20B**). The L98 residue's side chain is too short to contact the hydrophobic residues of the central helix of coGFP (**Figure S20A**).

The V162D is destabilizing the structures, probably due to the presence of residue E194 in a close proximity it and a potential electrostatic repulsion between these two equally charged side chains (**Table S2**). It is however possible, that pronounced destabilizing character of V162D mutation might be to some extent an artefact of the method, as the coGFP protein is negatively charged and every mutation increasing the negative charge, like V162D, is estimated as very unfavorable to the stability, whereas in reality, its local effects might not be that substantial. As a result, the potentially slightly destabilizing mutation of V162D is overestimated.

These results are in agreement with enrichments analysis (**Figure S15**) as the V162D mutation is observed frequently only under green fluorescence selection, otherwise this mutation is not frequent. On the other hand, the L98M mutation is being observed frequently in all populations under all selection regimes, thanks to its structurally stabilizing role.

When paired with V162D, the otherwise stabilizing G147S mutation has, surprisingly, destabilizing effect, despite the possibility of hydrogen bond formation with the aspartate residue 162. It is in agreement with the fraction of folded protein measured for V162D_G147S that was lowest from all of the measured proteins (**Figure S19**). However, in case of the triple mutant, V162D, G147S, L98M, the protein has a higher structural stability than the sum of the single mutations' ΔΔG values, pointing to synergistic effect of these mutations (**Table S2**) that is in agreement with the triple mutant's improved folding ability (**Figure S19**).

**Table S3. Detailed statistics for data reported in Fig. 5.** The test is based on a generalized linear model (binomial model, Mutation counts vs library type, likelihood ratio test). Beta means enrichment, p-values are bonferroni corrected. Negative beta means enriched in single-copy populations compared to double-copy populations. Positive beta means enriched in double-copy populations compared to single-copy populations.

| Mutation | Generation | Median frequency in single-copy populations | Median frequency in double-copy populations | p value | beta |
|---|---|---|---|---|---|
| G147S | 1 | 0.38 | 0.52 | 1.00E+00 | 0.38 |
| G147S | 2 | 0.51 | 2.01 | 2.21E-112 | 1.51 |
| G147S | 3 | 0.07 | 5.55 | 0.00E+00 | 3.95 |
| G147S | 4 | 7.89 | 1.26 | 1.00E+00 | -0.02 |
| G147S | 5 | 5.58 | 12.98 | 0.00E+00 | -1.81 |
| V162D | 1 | 0.11 | 0.30 | 1.49E-02 | 1.31 |
| V162D | 2 | 1.98 | 5.85 | 4.99E-244 | 1.06 |
| V162D | 3 | 92.13 | 89.02 | 1.26E-01 | 0.06 |
| V162D | 4 | 99.41 | 98.74 | 6.21E-12 | -1.20 |
| V162D | 5 | 98.99 | 99.23 | 4.25E-03 | -0.41 |
| L98M | 1 | 0.68 | 0.93 | 5.23E-01 | 0.53 |
| L98M | 2 | 4.90 | 4.50 | 1.00E+00 | -0.02 |
| L98M | 3 | 4.27 | 8.29 | 2.27E-122 | 0.58 |
| L98M | 4 | 5.44 | 20.00 | 4.28E-210 | 2.11 |
| L98M | 5 | 10.00 | 11.13 | 7.96E-138 | 0.59 |
| L98M+G147S | 1 | 0.00 | 0.00 | 1.00E+00 | 1.04 |
| L98M+G147S | 2 | 0.01 | 0.00 | 4.79E-02 | -1.65 |
| L98M+G147S | 3 | 0.00 | 0.00 | 9.42E-141 | 4.11 |
| L98M+G147S | 4 | 0.00 | 0.69 | 5.29E-08 | 2.51 |
| L98M+G147S | 5 | 0.64 | 4.24 | 0.00E+00 | -2.28 |
| L98M+V162D | 1 | 0.00 | 0.00 | 1.00E+00 | 1.04 |
| L98M+V162D | 2 | 0.00 | 0.00 | 1.00E+00 | -0.67 |
| L98M+V162D | 3 | 0.06 | 4.02 | 0.00E+00 | 3.22 |
| L98M+V162D | 4 | 5.08 | 19.43 | 2.91E-186 | 2.09 |
| L98M+V162D | 5 | 9.67 | 10.95 | 1.11E-114 | 0.54 |
| G147S+V162D | 1 | 0.00 | 0.00 | 1.00E+00 | 1.32 |
| G147S+V162D | 2 | 0.00 | 0.00 | 1.00E+00 | 0.25 |
| G147S+V162D | 3 | 0.00 | 0.00 | 1.00E+00 | -0.34 |
| G147S+V162D | 4 | 7.89 | 0.69 | 1.00E+00 | -0.16 |
| G147S+V162D | 5 | 5.58 | 12.91 | 0.00E+00 | -1.82 |
| L98M+G147S+V162D | 1 | 0.00 | 0.00 | 1.00E+00 | 1.04 |
| L98M+G147S+V162D | 2 | 0.00 | 0.00 | 1.00E+00 | 0.25 |
| L98M+G147S+V162D | 3 | 0.00 | 0.00 | 1.00E+00 | 0.17 |
| L98M+G147S+V162D | 4 | 0.00 | 0.57 | 8.99E-08 | 2.49 |
| L98M+G147S+V162D | 5 | 0.64 | 4.24 | 0.00E+00 | -2.28 |

**Table S4. Primers used in this study.**

| Oligonucleotide | Oligonucleotide sequence (5 → 3) |
|---|---|
| Seq_0_f | GAGTTGTAAAACGACGGCCAG |
| Seq_2_r | GAAAGCTGGTCCAAGCGATTG |
| Seq_3_f | CTCATTCGCTAATCGCCAC |
| pBAD_f | GCCGTCACTGCGTCTTTTAC |
| LJM01_f | GTG ATG ATG GTG ATG ATG GCC CAT ATG TAT ATC TCC |
| LJM01_r | GTG ATG ATG GTG ATG ATG GCC CAT GGT ATA TCT CCT |
| LJM02 | GAT ATA CAT ATG GGC CAT CAT CAC CAT CAT CAC AGC ATT CCG GAA AAT |
| LJM03_r | GTTACCAAACTGGAACCGGCGAGCGAAAGCATGTATGTTAG |
| LJM03_f | CTAACATACATGCTTTCGCTCGCCGGTTCCAGTTTGGTAAC |
| LJM04_f | GTTACCAAACTGGAACCGGGCAGCGAAAGCATGTATGTTAG |
| LJM04_r | CTAACATACATGCTTTCGCTGCCCGGTTCCAGTTTGGTAAC |
| LJM05 | GGC CAT CAT CAC CAT CAT CAC |
| LJM06_f | CTTATTCGGCCTTGAATTGATTATATGCGGATTAGAAAAACAACT |
| LJM06_r | AGTTGTTTTTCTAATCCGCATATAATCAATTCAAGGCCGAATAAG |
| LJM07_r | CAACTCGAATTCTTCCACCGTACGTCGAGCGGGAG |
| LJM08_f | GATATAGCGGCCGCAATGGCGGCGCGCCATCGAATG |
| LJM09_f | GTCATGGAATTCGAGTTGTAAAACGACG |
| LJM09_r | GATTATGCGGCCGCGCCGTCACTGCGTCTTTTAC |
| LJM10_f | GCTAGC CCATGG GCCATCATCATCACCATCATAG |
| LJM10_r | CTCTAC GGTACC TTATTACGGTTTGGCAATTGCGGTTTC |

**Table S5. List of site-directed mutagenesis primers.**

| Mutation | Forward primer | Reverse primer |
|---|---|---|
| Q74A, Y75S, G76A | GATATTCTGAGCGTTGCATTT GCC AGC GCG AATCGTACCTATACCAGCTATC | GATAGCTGGTATAGGTACGATT CGC GCT GGC AAATGCAACGCTCAGAATATC |
| V169D | GGTGAAGATGTTCTGAGCTATAAAACCCAG AGCACCCATT | CAGAACATCTTCACCAACCAGGGTGCCATCAC TAACATAC |
| P142L | GATGGTCTGGTTATGAAAAAAGAAGTTACC AAACTGGAAC | CATAACCAGACCATCTTCCGGGAAACCTTCAC CGTTATAT |
| Y173F | CTGAGCTTTAAAACCCAGAGCACCCATTATA CCTGTCACA | GGTTTTAAAGCTCAGAACAACTTCACCAACCA GGGTGCCA |
| S9R | CATCACCGCATTCCGGAAAATAGCGGTCTG ACCGAAGAAA | CGGAATGCGGAATGCTGTGATGATGGTGATGA TGGCCCAT |
| S9I | CATCACATCATTCCGGAAAATAGCGGTCTG ACCGAAGAAA | CGGAATGATGAATGCTGTGATGATGGTGATGA TGGCCCAT |
| S9C | CATCACTGCATTCCGGAAAATAGCGGTCTG ACCGAAGAAA | CGGAATGCAGAATGCTGTGATGATGGTGATGA TGGCCCAT |
| L105M | CGTACCATGAGCTTTGAAGATGGTGCCATTG TTAAAGTGG | AAAGCTCATGGTACGTTCAAAGGTAAAACCTT CCGGAAAG |
| G154A | GAACCGGCCAGCGAAAGCATGTATGTTAGT GATGGCACCC | TTCGCTGGCCGGTTCCAGTTTGGTAACTTCTTT TTTCATA |
| G154C | GAACCGTGCAGCGAAAGCATGTATGTTAGT GATGGCACCC | TTCGCTGCACGGTTCCAGTTTGGTAACTTCTTT TTTCATA |
| G154S | GAACCGAGCAGCGAAAGCATGTATGTTAGT GATGGCACCC | TTCGCTGCTCGGTTCCAGTTTGGTAACTTCTTTT TTCATA |
| G154D | GAACCGGACAGCGAAAGCATGTATGTTAGT GATGGCACCC | TTCGCTGTCCGGTTCCAGTTTGGTAACTTCTTTT TTCATA |
| G48S | CTGACCAGTATTCAGAAACTGGATATTCGTG TTATTGAAG | CTGAATACTGGTCAGAATATTACCACCACCAA TACCTTCC |
| T79N | AATCGTAACTATACCAGCTATCCGGCAAAA ATCCCGGATT | GGTATAGTTACGATTGCCATACTGAAATGCAA CGCTCAGA |
| V127L | AAATTTCTGGGCAAAATCAAATATAACGGT GAAGGTTTCC | TTTGCCCAGAAATTTACCATCCTCGATGCTAAT ATCGCTT |
| G76D | CAGTATGACAATCGTACCTATACCAGCTATC CGGCAAAAA | ACGATTGTCATACTGAAATGCAACGCTCAGAA TATCAAAG |
| G42A | ATTGGTGCTGGTAATATTCTGACCGGTATTC AGAAACTGG | ATTACCAGCACCAATACCTTCCATGCTAAAGG CATGACCA |
| H183R | ACCTGTCGCATGAAAACCATTTATCGCAGCA AAAAACCGG | TTTCATGCGACAGGTATAATGGGTGCTCTGGGT TTTATAG |
| K129R | GTGGGCAGAATCAAATATAACGGTGAAGGT | TTTGATTCTGCCCACAAATTTACCATCCTCGAT |

| | | |
|---|---|---|
| | TTCCCGGAAG | GCTAATA |
| S155I | CCGGGCATCGAAAGCATGTATGTTAGTGATGGCACCCTGG | GCTTTCGATGCCCGGTTCCAGTTTGGTAACTTCTTTTTTC |
| E168D | GTTGGTGATGTTGTTCTGAGCTATAAAACCCAGAGCACCC | AACAACATCACCAACCAGGGTGCCATCACTAACATACATG |
| G163D | AGTGATGACACCCTGGTTGGTGAAGTTGTTCTGAGCTATA | CAGGGTGTCATCACTAACATACATGCTTTCGCTGCCCGGT |
| L197M | GAAAACATGCCGAAATTTCATTATGTTCATCACCGCCTGG | TTTCGGCATGTTTTCAACCGGTTTTTGCTGCGATAAATG |
| O229Y | AAACCGTATTAAGAGCTCCAATCGCTTGGACCAGCTTTCC | CTCTTAATACGGTTTGGCAATTGCGGTTTCATGCTGCTCG |
| R206L | CATCACCTCCTGGAAAAAAAAATTGTGGAAGAGGGCTATT | TTCCAGGAGGTGATGAACATAATGAAATTTCGGCAGGTTT |
| S106N | ACCCTGAACTTTGAAGATGGTGCCATTGTTAAAGTGGAAA | TTCAAAGTTCAGGGTACGTTCAAAGGTAAAACCTTCCGGA |
| S177N | ACCCAGAACACCCATTATACCTGTCACATGAAAACCATTT | ATGGGTGTTCTGGGTTTTATAGCTCAGAACAACTTCACCA |
| S155N | CCGGGCAACGAAAGCATGTATGTTAGTGATGGCACCCTGG | GCTTTCGTTGCCCGGTTCCAGTTTGGTAACTTCTTTTTTC |
| S172R | GTTCTGAGATATAAAACCCAGAGCACCCATTATACCTGTC | TTTATATCTCAGAACAACTTCACCAACCAGGGTGCCATCA |
| V148I | AAAGAAATTACCAAACTGGAACCGGGCAGCGAAAGCATGT | TTTGGTAATTTCTTTTTCATAACCGGACCATCTTCCGGG |
| R206H | CATCACCACCTGGAAAAAAAAATTGTGGAAGAGGGCTATT | TTCCAGGTGGTGATGAACATAATGAAATTTCGGCAGGTTT |
| H183R | ACCTGTCGCATGAAAACCATTTATCGCAGCAAAAAACCGG | TTTCATGCGACAGGTATAATGGGTGCTCTGGGTTTTATAG |

**Table S6. Plasmids used in this study.**

| Plasmid name | Description | Source | Addgene number |
|---|---|---|---|
| pAND | Source of backbone | Addgene #49377 | #49377 |
| pAND-MCS | MCS added, NdeI site removed from TetR | This study | #223514 |
| pDUP | *cogfp* under $P_{tac}$ | This study | #223515 |
| pDUP1 | *cogfp* inactive under $P_{tet}$, *cogfp* under $P_{tac}$ | This study | #223516 |
| pDUP2 | *cogfp* under $P_{tet}$, *cogfp* under $P_{tac}$ | This study | #223517 |
